# Dense reconstruction of brain-wide neuronal population close to the ground truth

**DOI:** 10.1101/223834

**Authors:** Zhou Hang, Li Shiwei, Li Anan, Xiong Feng, Li Ning, Han Jiacheng, Kang Hongtao, Chen Yijun, Li Yun, Fang Wenqian, Liu Yidong, Lin Huimin, Jin Sen, Li Zhiming, Xu Fuqiang, Zhang Yu-hui, Lv Xiaohua, Liu Xiuli, Gong Hui, Luo Qingming, Quan Tingwei, Zeng Shaoqun

## Abstract

Recent progresses allow imaging specific neuronal populations at single-axon level across mouse brain. However, digital reconstruction of neurons in large dataset requires months of human labor. Here, we developed a tool to solve this problem. Our tool offers a special error-screening system for fast localization of submicron errors in densely packed neurites and along long projection across the whole brain, thus achieving reconstruction close to the ground-truth. Moreover, our tool equips algorithms that significantly reduce intensive manual interferences and achieve high-level automation, with speed 5 times faster compared to semi-automatic tools. We also demonstrated reconstruction of 35 long projection neurons around one injection site of a mouse brain at an affordable time cost. Our tool is applicable with datasets of 10 TB or higher from various light microscopy, and provides a starting point for the reconstruction of neuronal population for neuroscience studies at a single-cell level.

## Introduction

Mapping neuronal morphology at single-cell level will bridge the gap between micro scale and macro scale studies^1-3^, and play an important role in cell type, neural circuits, and neural computing studies^4-6^. Recent breakthroughs in imaging^7-10^ and molecular labeling ^11, 12^ techniques have provided Terabytes (TBs)-sized dataset from which we can measure almost the complete morphology of the neuronal population at a single-axon resolution (**Video 1**). Special data format and data splitting mode have been developed to browse or visualize TB^13, 14^ or PB (Petabytes)^15^ dataset. However, tracing these brain-wide neuronal projections is challenging^16, 17^ Automatic tracing tools ^18-22^ applicable to local neuronal population (Gigabytes size) cannot extend to whole brain. Just like “snowball effect”, for a neuron with tree-like structure, a micron-sized error in tracing will result in a huge error^23^, as subsequent tracing is not credible and may extend to the whole brain (**Supplementary Fig. 1**). The densely packed neurite makes this situation even worse, as dense neurite tracing is still an open question^1, 17, 23, 24^, usually leads to lower tracing accuracy. Practically semi-automatic tools^25, 26^ have been applied for neuronal tracing in large dataset and its precision is well acknowledged. However, it is extremely laborious and time-consuming for brain-wide reconstruction, as these tools usually work at very low automatic level. Reconstructing a population of neurons at brain-wide scale will require thousands of hours of manual labors.

**Figure 1.**
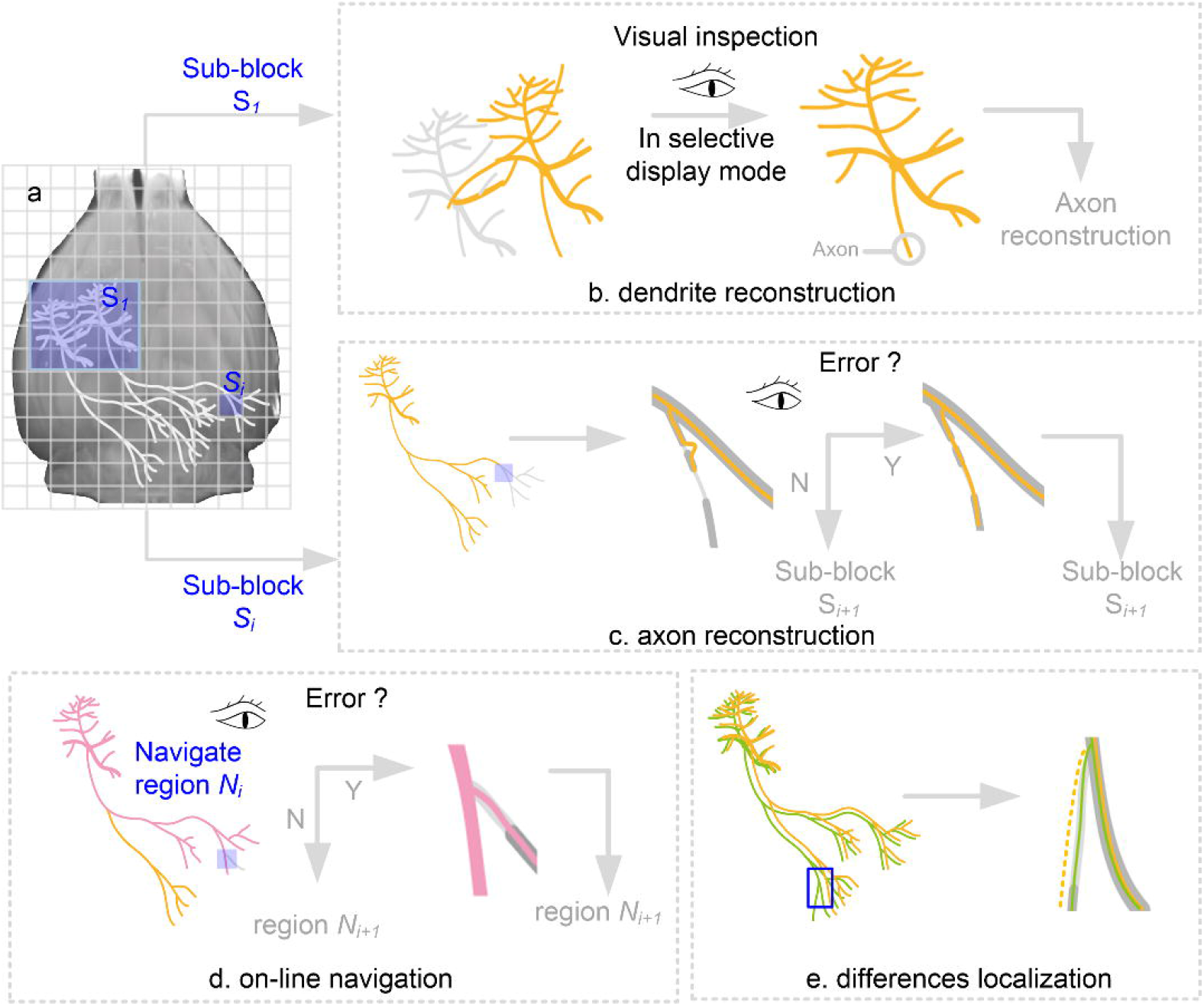
The pipeline of brain-wide reconstruction of neuronal population. (a) Divide the whole brain images into sub-blocks with the same size, and select the 3D region of interest (the 1st ROI) according to the known position of a soma that corresponds to the target neuron. ROI refers to the interested dataset including some divided sub-blocks. (b) Reconstruct the target neuron and other neurons in the1st ROI. Revise the reconstruction of the target neuron in selective display mode, and find the initial part of the axon for axonal tracing. (c) Trace axons in the current ROI and revise the traced results; determine the new ROI that contains the locations where traced axons touch the boundary of the current ROI. (d) Reconstruction navigation for spotting reconstruction errors at brain-wide scale. (e) Online locate the differences between the reconstructions of a neuron performed by two annotators, and check the reconstruction differences for trustworthy reconstruction.

Here, we built a global tree reconstruction system (GTree) for brain-wide population reconstruction. In GTree, we developed a selective display mode to boost local dense reconstruction and acquired a higher accuracy than the well-acknowledged reconstruction accuracy generated with semi-automatic software. Moreover, we introduced brain-wide error-screening system to improve the precision across whole brain. Furthermore, we introduced algorithms that significantly reduce intensive manual interferences and thus achieve high-level automated reconstruction. With all these efforts, we demonstrated a brain-wide population reconstruction, with precision close to ground-truth, speed at least 5 times faster compared to semi-automatic tools. We also demonstrated a successful reconstruction of 35 long projection neurons around one injection site of a mouse brain at an affordable time cost.

## Design of GTree

In addition to local neuronal population reconstruction, GTree can achieve brain-wide reconstruction (**Online Methods**). In the design of GTree for brain-wide reconstruction, we built functions corresponding to key reconstruction steps, which included dendrite reconstructions, axon reconstructions and checking the reconstructions on a brain-wide scale (**Fig. 1**). In addition, a technique for multi-resolution representation of a TB-sized dataset is necessary (**Supplementary Note 1**).

We divided the whole-brain dataset into many sub-blocks with a multi-resolution representation technique^15^ and selected the first sub-block (e.g., a 3D region of interest (ROI) including the target neuron (soma)) to begin tracing (**Fig. 1a**). In general, the first ROI mainly contains somas and dendrites. In the first ROI, we used NeuroGPS-Tree^18^ to automatically reconstruct the target neuron and other neurons. We revised the automated reconstruction of the target neuron in a selective display mode (**Fig. 1b**) in which reconstruction errors can be easily found. GTree can automatically record the locations where traced dendrites touch the boundary of the first ROI simultaneously. These locations provide cues to select the subsequent ROIs to trace the dendrites of the target neuron.

After completing the reconstruction of dendrites, we located the initial part of the axon for axonal tracing. Similar to the tracing of dendrites, a human-supervised checking procedure was implemented for trustworthy reconstruction (**Fig. 1c**). Note that except for the first ROI, in which packed neurites are usually included, the neurites in all ROIs are sparsely distributed in our analysis. Therefore, we traced neurites in the first ROI with NeuroGPS-Tree and others with SparseTracer^27^, as SparseTracer presents a faster reconstruction speed than NeuroGPS-Tree when neurites are sparsely distributed.

As described for the above procedures (**Figs. 1b-c**), human-supervised reconstruction is performed in every sub-block. Brain-wide reconstruction of a neuron requires the analysis of hundreds of sub-blocks, and reconstruction errors are still unavoidable. GTree uses reconstruction navigation (**Supplementary Note 2**) to identify reconstruction errors on a brain-wide scale (**Fig. 1d**). With this navigation, we can vastly browse the reconstruction and the neighboring image region. The browsing information is used for checking the reconstructions. By using this navigation module, one can vastly check a brain-wide reconstruction within 1.5-2 hours. Furthermore, for the generation of a ground-truth reconstruction, the general strategy is that different annotators reconstruct the same neuron and regard the common consent as the ground truth ^28^. In this process, locating the difference between the reconstructions (**Fig. 1e**) is an indispensable step. Based on hash query method (**Supplementary Note 3**), GTree locates differences between brain-wide reconstructions within a second, which is several hundred times faster than the common way (**Supplementary Fig. 2**).

**Figure 2.**
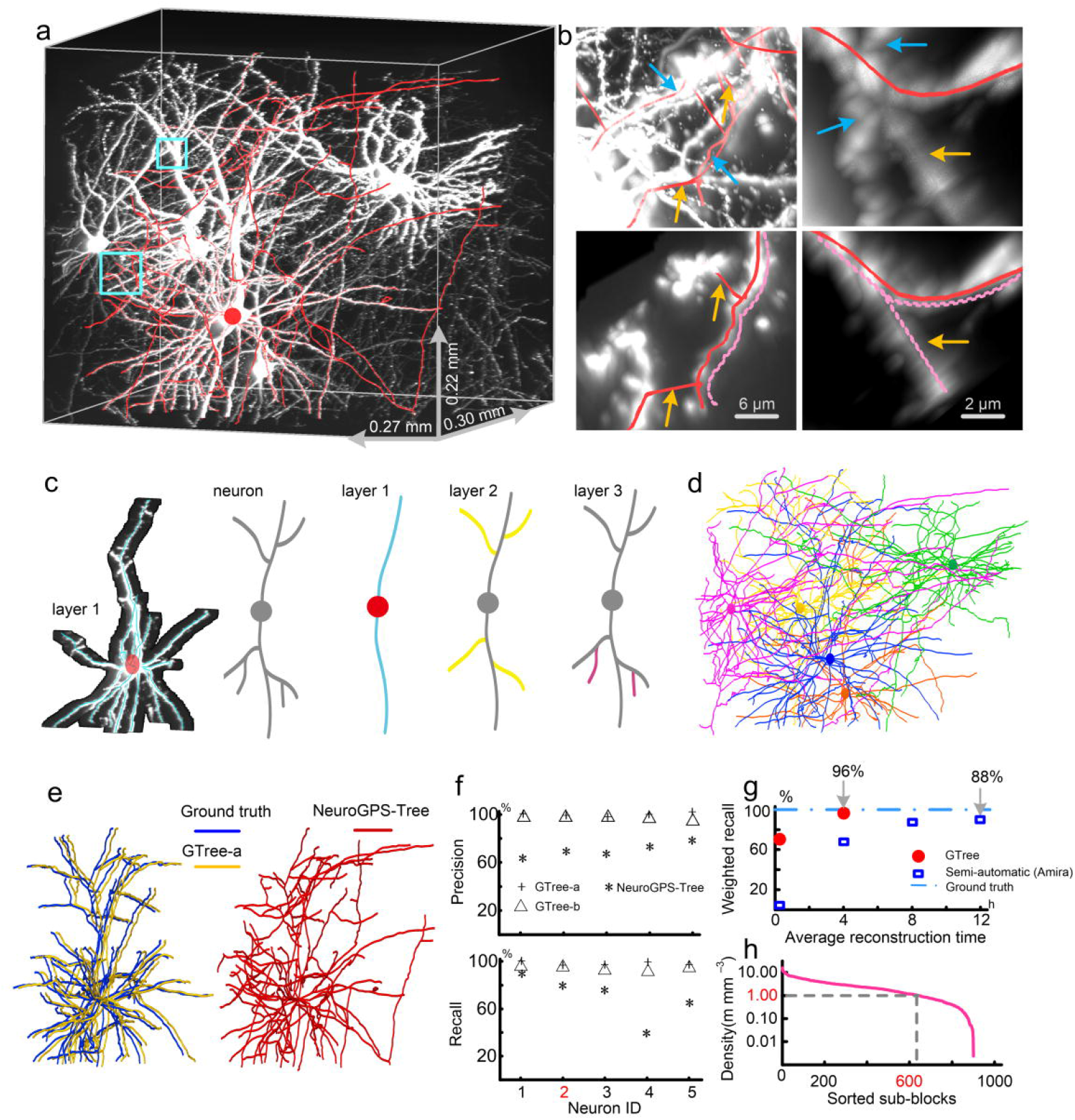
GTree achieved a dense reconstruction with high precision. (a) The imaging dataset from the neocortex had a size of 1343 × 1120 × 302 voxels and contained packed dendrites and axons. The red curve represents the reconstruction of a neuron generated with the automatic method. (b) Reconstruction errors are shown without (upper panels) and with (bottom panels) the selective display mode and are labeled with orange arrows. The neurites labeled with blue arrows (upper panels) seriously hinder error checks and were removed in the selective display mode. The dashed and red lines represent the ground-truth reconstruction and the automated reconstruction, respectively (bottom panels). (c) For the reconstructed neuron (red in (a)), the reconstruction of neurites in layer 1 (blue curves) and their neighboring image regions are displayed (left). This display is based on the tree-like structure in which neurites can be assigned to their corresponding layer (right). (d) A population of five neurons was reconstructed from (a), and the neurons are displayed with different colors. (e) Comparison of the reconstruction of a neuron with GTree and the ground-truth reconstruction. The automated reconstruction with NeuroGPS-Tree is also presented. (f) Quantification of the reconstructions driven by GTree and NeuroGPS-Tree with recall and precision rates. GTree-a and -b refer to the reconstructions of the same population performed by two groups of skilled annotators. (g) Comparison of the reconstruction performance of GTree and semi-automatic software (Amira). (h) Spatial neurite density distribution, obtained by calculating the total length of neurites in a sub-block with a size of 100 × 100 × 25 voxels (10^−5^ mm^3^).

Based on the entire process of reconstruction, we demonstrated that, in addition to the selective display mode (**Online Methods**), GTree offers a system for thoroughly checking reconstruction errors on a brain-wide scale (**Online Methods**). This system runs through the pipeline of reconstruction and assures a high-precision reconstruction.

## Dense reconstruction

We evaluated the dense reconstruction performance of GTree using a typical dataset from the neocortex. The dataset contained packed neurites with a wide range of signal intensities (**Fig. 2a** and **Video2**) and therefore challenged current automatic reconstruction methods (**Supplementary Fig. 3**). For highly accurate dense reconstruction, we designed the selective display mode in GTree (**Online Methods**). This mode enabled us to focus on checking the target reconstructions and their related signals (**Fig. 2b**) by removing other unrelated signals. Without the selective display mode, it is time consuming and laborious for a skilled annotator to find reconstruction errors due to interference from other signals (upper panels in **Fig. 2b**). The selective display mode eliminated this kind of interference and allowed effective checking for reconstruction errors (bottom panels in **Fig. 2b**). In the design of the selective display mode, we considered the fact that the reconstructed neurons could be mapped as a tree graph in which a node represents a traced neurite (**Supplementary Fig. 4**). Due to this mapping, the corresponding reconstructions and their neighborhood images can be visualized when selecting the sub-structure of the tree graph (**Fig. 2c**). In the selective display mode, we revised the automated reconstruction from the population dataset (**Fig. 2a**) generated with NeuroGPS-Tree. We present the revised reconstruction in which an individual neuron is displayed with a different color (**Fig. 2d** and **Video 2**). To quantify the performance of the reconstructions driven by GTree and semi-automatic software, we designed a procedure for reducing reconstruction variance among different annotators. Two annotators were classified into one group. One presented the reconstructions, and the other checked the presented reconstructions. The total time was set to 4 hours (a third of the time was occupied by checking reconstructions) or 12 hours per neuron for GTree and the semi-automatic software (Amira^26^), respectively. Using this procedure, GTree produced two reconstructions of the same population (GTree-a and -b). We present the reconstruction of one neuron from GTree-a that exhibited obvious differences from the reconstruction provided by NeuroGPS-Tree but was close to the ground-truth reconstruction (**Fig. 2e**).

**Figure 3.**
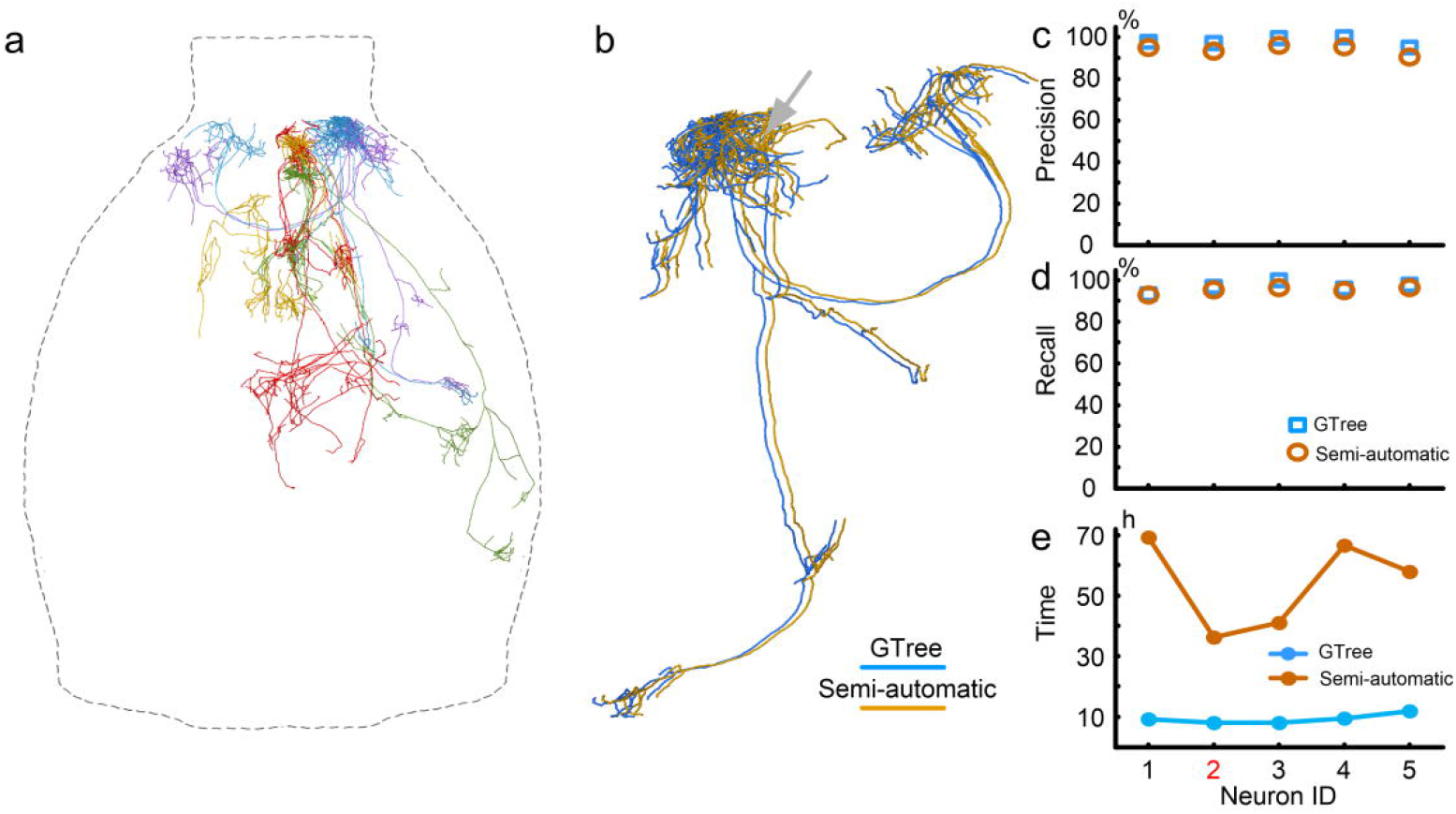
High-precision reconstruction on a brain-wide scale with GTree. (a) Reconstruction of five neurons using GTree. (b) Comparison of the reconstructions of a neuron derived from GTree and semi-automatic software. A difference (gray arrow) between the two reconstructions is shown. Quantification of the reconstructions based on precision (c) and recall rates (d). (e) Comparison of reconstruction time costs using GTree and semi-automatic software.

**Figure 4.**
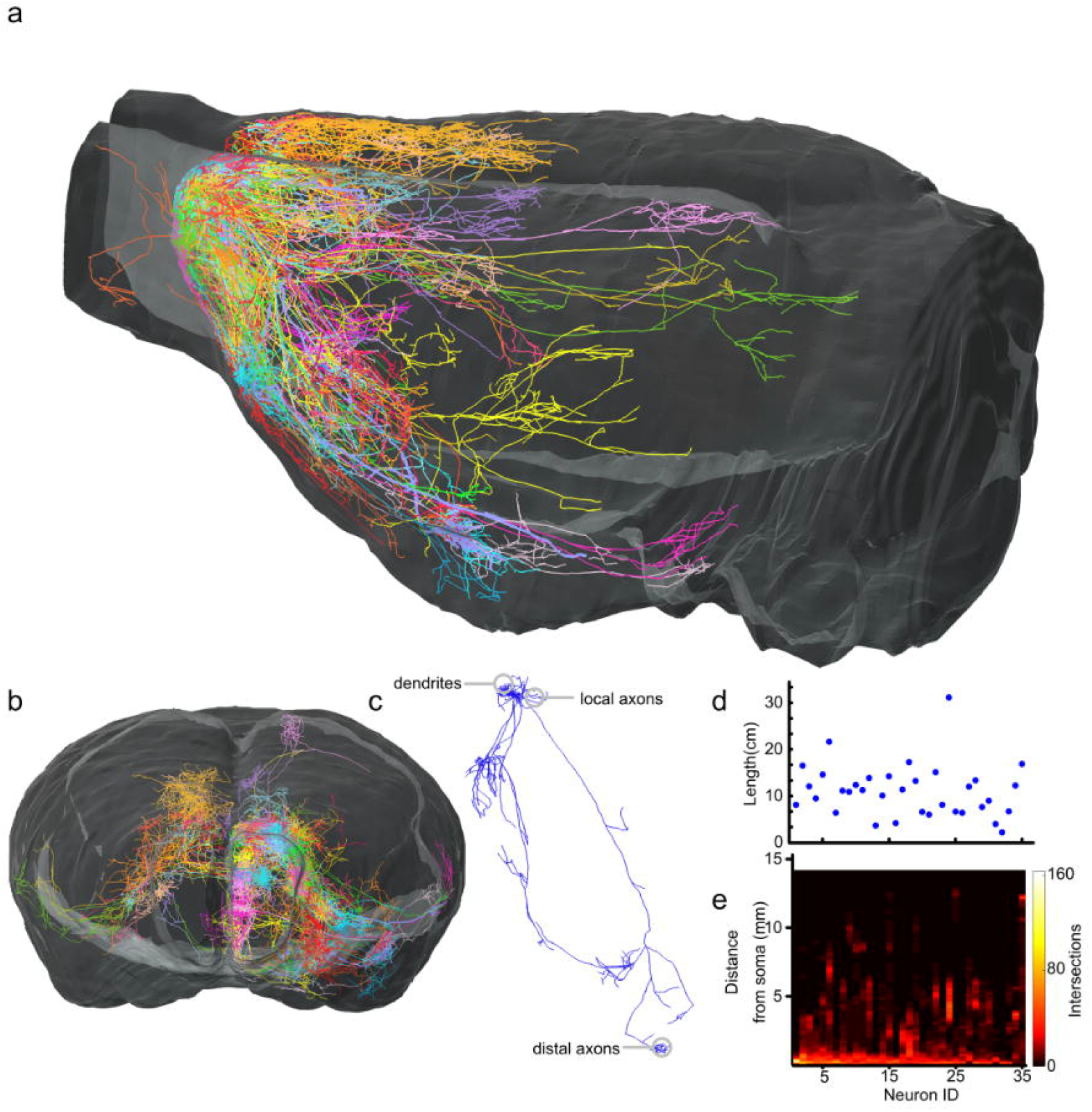
Reconstruction of the neuronal population on a brain-wide scale. (a) The neuronal population across the whole mouse brain, including 35 neurons, was reconstructed. The imaging dataset was 10 TBs in size. The individual reconstructed neurons are identified using different colors. (b) The reconstructions are displayed in the coronal view. (c) A reconstructed neuron, including dendrites, local axons and distal axons. (d) The total length of neurites identified in the individual reconstructed neuron. (e) Sholl analysis of reconstructed neurons. In this analysis, a series of sphere surfaces are generated in 20 μm radial increments centered on the neuronal soma, and the reconstructed neurites that intersect each sphere surface are then counted. This operation constructed the relationships between the number of intersections and distances from the soma.

We quantified the reconstructions from the population dataset (**Fig. 2a**), including 5 neurons with recall and precision rates (**Fig. 2f**). To calculate these two evaluation indexes, ground-truth reconstruction is necessary and can be achieved via the strategy employing voting to reach a consensus on the discrepancies among the reconstructions performed by three annotators^28^. We found that GTree could provide higher reconstruction accuracy than NeuroGPS-Tree (**Fig. 2f**). The recall and precision rates were 96% and 97%, respectively, for GTree (GTree-a) versus 70% and 69%, respectively, for our previously developed software, NeuroGPS-Tree. This difference in reconstruction was significant. We also found that there were no significant differences between the two groups of reconstructions generated with GTree (Kolmogorov–Smirnov test, precision: p > 0.2; recall: p > 0.69), indicating that GTree can provide robust reconstructions.

Furthermore, we demonstrated that GTree provided more accurate reconstruction than the semi-automatic software, which shows a widely accepted high accuracy. Considering a few false reconstructions, we used weighted recall to quantify the population reconstructions driven by GTree and semi-automatic software. Weighted recall refers to the weighted average recall rate of the reconstructions of individual neurons. The weight was proportional to the total length of the neurites of an individual neuron identified through ground-truth reconstruction. GTree reconstructed the neuronal population with an integral degree of 96% (**Fig. 2g**). The time cost was within 4 hours per neuron. In contrast, the semi-automatic software provided reconstruction accuracy with an integral degree of 88%, and the time cost was set to 12 hours per neuron. In addition, increasing the reconstruction time (from 8 to 12 h) did not boost reconstruction accuracy (**Fig. 2g**). This phenomenon demonstrates that it is an extremely difficult task for semi-automatic software to even slightly improve the accuracy for a population dataset with massively packed neurites (**Fig. 2h**).

## High-precision reconstruction on a brain-wide scale

We evaluated performance of GTree in brain-wide reconstruction with the 10 TBs-sized images. The dataset contained specially labeled neurons whose morphologies spanned different brain regions or even the whole brain. This imaging dataset provide sufficient information for trustworthy reconstruction including dendrites and distal axons. We randomly selected five neurons in this dataset and generated their ground-truth reconstruction (**Online Methods**) to evaluate reconstruction performance between GTree and the semi-automatic software. We present the reconstructions of these five neurons obtained with GTree (**Fig. 3a**). The results indicated that the reconstructed neurons exhibited many neurites and that these neurites were distributed in different brain regions. We also used semi-automatic tool (Amira) to manually reconstruct these five neurons. When we compared the semi-automatic reconstruction of a neuron with the reconstruction provided by GTree, we found that the differences between these two reconstructions were negligible (**Fig. 3b & Supplementary Fig. 5**). We further quantified the reconstructions of these five neurons based on recall and precision rates (**Figs. 3c & d**). The results showed that the weighted average recall and precision rates were 96% and 98%, respectively, for GTree, versus 95% and 94% for the semi-automatic software. The weights were proportional to the total length of neurites identified in the ground-truth reconstruction. The reconstruction results were similar between GTree and semi-automatic software. We concluded that GTree, like the semi-automatic software, can provide a brain-wide reconstruction close to the ground truth. Furthermore, we quantified the time costs of the reconstruction of these five neurons using GTree and semi-automatic software. The results showed that GTree spent approximately 9 hours on the reconstruction of a single neuron, which was at least five times faster than the semi-automatic software (**Fig. 3e**). In this comparison, we selected annotators who were well trained (**Online Methods**), and these annotators acquired the reconstruction on the same computing platform equipment with Redundant Arrays of Independent Disks (RAID).

## High-level automatic reconstruction

In GTree, we developed a series of algorithms for those reconstruction steps that required intensive manual labor. Two algorithms are provided here as examples. One is used for detecting the optimal skeleton of a neurite, and the other is used for identifying the neurites with weak signals. We based on a Lasso model^29^ to correct the reconstructed skeleton points, including bifurcation points, to their optimal positions. The optimal positions are generally located at the centerline of a neurite and exhibit the maximum signal intensities. We evaluated the Lasso-based model using the dataset of axons (Supplementary Fig. 6a). In the reconstruction process, there were 19 sub-blocks in which the reconstructed skeletons of the tortuous neurites had to be revised. With the Lasso model, the reconstructed skeletons in 17 sub-blocks were automatically corrected, avoiding the corresponding human editing. We present two typical detection examples (**Supplementary Figs. 6b-e**) showing the effectiveness of the Lasso-based model. We also built a machine-learning method for identifying neurites with weak signals^30^. We demonstrated that the identification method could vastly reduce the human editing of the reconstructions using 3 sub-blocks of datasets (**Supplementary Fig. 7a**). In the analysis of these selected datasets, when the identification method is equipped, the total number of human edits was reduced to 18, versus 136 without the identification method (**Supplementary Figs. 7b-d**). In addition, we calculated the signal-to-background ratios of the points in the reconstructed skeletons (**Supplementary Fig. 7e**). The points with SBR < 1.3 occupied 80% percent of all skeleton points, which can explain why a large amount of human editing is required without the use of the identification method.

## Brain-wide population reconstruction

We used GTree to reconstruct a population of neurons from the whole mouse brain imaging dataset. The imaging dataset included 35 pyramidal neurons which are from one injection site. Despite the sparse labeling, packed neurites were still commonly found in the imaging dataset. Using GTree, all of the labeled neurons in this imaging dataset could be thoroughly reconstructed (**Figs. 4a &b, Video 3, Supplementary Fig. 8**). A typical neuron could be nearly completely reconstructed, such that dendrites, local axons, and distal axons can be easily identified (**Fig. 4c**). The reconstructions allowed the application of quantitative morphological measures, such as the total length of neurites per neuron (**Fig. 4d**) and Sholl analysis^31^ (**Fig. 4e**). According to the information on the reconstruction speed (**Fig. 3e**) and the measurement of the reconstructions (**Fig. 4d**), we roughly estimated that reconstructing all 35 of these neurons with semi-automatic software would require more than 1300 hours (total length of neurites, 396.7 cm; speed, 3.4 hours per centimeter). GTree reduced the time cost to less than 250 hours (speed, 0.6 hours per centimeter). These results indicated that extremely heavy labor is required for brain-wide population reconstruction with semi-automatic software, and this current reconstruction status can be changed using GTree.

## Applicability

In addition to the fMOST datasets, we demonstrated the applicability of GTree to other types of datasets. These datasets were collected with different imaging modalities, such as serial-two-photon tomography^8^, light-sheet^32-34^, two-photon microscopy^35^ and so on. The axial resolution of an STP dataset is low, at 10 μm. In this case, GTree reconstructed all four labeled neurons spanning different brain regions. The reconstructions were consistent with the ground truth^36^ (**Supplementary Fig. 9**). We further used GTree to reconstruct 5 groups of datasets collected via light-sheet microscopy^33^ (**Supplementary Fig. 10 & Supplementary Fig. 11**). The results suggested that GTree could provide a high-precision reconstruction from a dataset with a low optical spatial resolution. In addition, two types of public datasets, the Diadem and BigNeuron datasets acquired using wide-field, confocal, or two-photon microscopy, were used to test GTree. In the Diadem datasets^37^, we selected Neocortical Layer 1 Axons (**Supplementary Fig. 12**) and Hippocampal CA3 Interneuron image stacks (**Supplementary Fig. 13**) and used GTree to analyze them. The reconstructions showed that GTree behaved well in dense reconstructions and in the presence of extremely weak signals. Using BigNeuron datasets ^6, 38^, we demonstrated that GTree could detect weak signals in the presence of strong noise points (**Supplementary Fig. 14**) and could also detect torturous neurites (**Supplementary Fig. 15**). From the above testing results, we concluded that GTree can be widely used in neuron reconstruction.

## Discussion

In this study, we built the software tool GTree to achieve a dense reconstruction of the brain-wide neuronal population that challenges semi-automatic reconstruction with semi-automatic software. GTree achieved brain-wide reconstruction of neuronal populations with a speed gain of at least five times over semi-automatic software. Besides encompassed our established work, such as multiresolution representation of a dataset^15^, neurite tracing^27^, automatic reconstruction of a local neuronal population^18^, and identification of weak signals^30^, in GTree, we designed a suite of new techniques to overcome a series of challenges in brain-wide reconstruction, including the identification of packed neurites, checking for reconstruction errors in a TB-sized dataset, and improvement of automatic levels of reconstruction. Furthermore, GTree offers a system to check the reconstructions on different scales, from a local neurite to an entire single neuron. Due to these multitudinous and indispensable functional modules, GTree is an effective tool for brain-wide population reconstruction and may be helpful for many purposes, such as the identification of neuron types, investigation of the projection pattern of neurons, and mapping the neuronal circuitry.

Accurate brain-wide reconstruction of neuronal populations requires a series of image processing technologies, which can be integrated into a target software tool. As described previously, GTree has this characteristic and can therefore achieve brain-wide reconstruction with a high precision and relatively high throughput. This high precision can be attributed to the following factors. First, the selected display mode of dense reconstructions largely eliminates the interference from other reconstructions and can therefore effectively check the reconstructions. Second, online feedback for reconstructions from sub-blocks of brain images and editing function modules are combined to achieve supervised reconstructions. Third, fast navigation of the reconstructions and locating the differences in the reconstructions enable us to check the reconstructions on a brain-wide scale. There are two primary reasons for the relatively high-throughput reconstructions: one is the contributions of many automatic algorithms, including algorithms for neurite tracing^27^, spurious links between neurite identification^18^, weak signal identification^30^, and optimal skeleton detection ^29^, among many others; and the other is the user-friendly visualization and editing functions, which allows fast revision of reconstruction errors.

Dense reconstructions must essentially resolve closely spaced neurites, which is one of the most challenging problems in neuronal population reconstruction^1, 17, 23, 24^. To overcome this challenge, our previously developed automatic tool, NeuroGPS-Tree^18^, partially mimicked human strategies to identify individual packed neurites and achieved dense population reconstruction with an approximately 80% reconstruction accuracy. For a more challenging dataset packed with massive axons (**Fig. 2a**), the reconstruction accuracy was further decreased (**Fig. 2f**). This reconstruction accuracy was not sufficient for neuroscience research in many cases^4, 17^. In addition, the noted reconstruction accuracy was generated from the analysis of a GB-sized dataset. When NeuroGPS-Tree^18^ was extended to a TB-sized dataset, the corresponding reconstruction is not acceptable due to its fully automatic character (**Supplementary Fig. 1**). Considering this situation, we developed GTree for dense reconstruction. GTree integrates NeruoGPS-Tree and a selective display mode, thus effectively checking reconstructions provided by NeuroGPS-Tree. GTree vastly boosts the capability of NeuroGPS-Tree in dense reconstruction. GTree can generate dense reconstructions close to the ground truth (**Fig. 2f**) and therefore presents more applications. It should be noted that the selective display mode in GTree requires an initial population reconstruction. At present, NeuroGPS-Tree is a suitable tool for this purpose.

In GTree, the size of the dataset and the data format are essentially not restricted. The current version of GTree supports the tagged image file format (TIFF, 8/16 bits). Other data formats can be freely and easily converted into TIFF using third-party software such as ImageJ^39^. The data organization method (**Supplementary Note 1**) based on HDF5 was developed and integrated into GTree. Therefore, GTree can reconstruct neurons from hundreds of GBs sized dataset, without the help of other software. When the analyzed dataset increases to TBs or greater in size, this large data requires to be transformed into the big data format including TDat^15^. GTree supports the TDat format and thus can be suitable for reconstruction on TBs sized dataset. We also noted that other tools like Terafly^14^ and BigDataViewer^13^ perform well in the big data organization. However, these two tools have no related plugins for other software tools at present. So, the data formats generated with these two tools can’t be used in GTree.

The tree-like structure of neurons means that reconstruction error exerts a highly correlated effect on subsequent reconstructions. Without human supervision, one false reconstruction will lead to missing neurites or miss-assigned neurites, which explains why semi-automatic reconstruction is still the primary method for quantifying neurons^17^, despite the existence of numerous automatic algorithms. Whole-brain images are even more challenging, and human supervision is therefore necessary to obtain trustworthy results. However, the available semi-automatic reconstruction software lacks some key algorithms and editing functions to accurately identify an error in the presence of interference from densely paced neurites and cannot locate local reconstruction errors in centimeter-long projections^4, 17^. Therefore, semi-automatic reconstruction becomes time-consuming and laborious, making it difficult to match the development of neuronal images^16^. GTree fills this gap to a certain extent. In the near future, the level of automatic reconstruction by GTree will be further enhanced by introducing vast IO formats and automatic identification of potential errors.

## Online Methods

### 1. A brief description of the software tool

The GTree software is written in C++ and is freely available for academic research (**https://github.com/artzers/AdTree**). GTree can be divided into three components according to its functions: 1) reconstruction of a local neuronal population; 2) reconstruction of a brain-wide population; and 3) checking of reconstructions on a brain-wide scale.

#### Reconstruction of a local neuronal population

In this mode of reconstruction, the supported format of an input image is an 8 or 16 bit TIFF series (gray image stacks), and the output results are swc files^40^, which include the positions of skeleton points and the connections between the points. This component integrates the selective display mode and the editing function into our previously developed software tool NeuroGPS-Tree^18^. This integration greatly extends the ability of NeuroGPS-Tree and can provide a dense reconstruction close to the ground truth.

#### Reconstruction of a brain-wide neuronal population (Video 4)

Brain-wide reconstruction involves the analysis of TB-sized datasets. Therefore, a multi-resolution representative of a TB-sized dataset is required. Here, we convert a whole-brain dataset into a big data format, TDat^15^, in which the sub-blocks can be effectively loaded into computer memory. The reconstruction of a brain-wide neuronal population consists of reconstructions from a series of sub-blocks. SparseTracer^27^ is used for reconstructing neurites in sub-blocks. The corresponding editing functions are also matched for correcting reconstruction errors.

#### Checking reconstructions at brain-wide scale

Despite the editing function allowing checking of reconstructions from sub-blocks, brain-wide reconstruction is a long reconstruction process and includes the analysis of hundreds of sub-blocks. Thus, reconstruction errors are unavoidable. GTree provides two functions for checking reconstructions. One is reconstruction navigation on a brain-wide scale (Video 5); the other is localization of differences between reconstructions of the same neuron (Video 6). Reconstruction navigation can browse the reconstructed skeletons and their neighboring regions. The browsing information is used for checking reconstructions. The other function is to locate differences between different reconstructions of the same neuron. This function is necessary for high-precision reconstruction from a challenging dataset and ground-truth reconstruction.

### 2. Selective display mode for dense reconstruction

This mode is used for checking errors in dense reconstruction. When an automated method reconstructs a neuronal population with packed neurites, reconstruction errors are unavoidable and difficult to find because of the interference from other neurites, especially those that are closely positioned. In fact, a neuron can be mapped to a tree-graph in which a node and the connections between nodes represent a neurite and the links between neurites^18^, respectively (**Fig. 2c & Video** 7). Accordingly, the reconstructed neuron has the same map, meaning that when the sub-structure of this mapping graph is selected, the corresponding reconstruction (i.e., the reconstructed skeletons of neurites) is confirmed. Centered on the reconstructed skeleton, the cylindrical neighborhood regions are extracted from the original image. The radius of the neighborhood region is manually set. The reconstructions and the corresponding neighborhood images remain simultaneously and black out all of the signals that are not in the current regions. Using this operation, we can display the cylindrical region and the included reconstructions without interference from other signals. This display method is named after the selective display mode. In the selective display mode, we can focus on the interesting reconstructions and check them (**Video 8**). We recommend that checking is performed in the selective display mode (**Video 9**) to ensure that every neurite in the reconstruction is checked.

### 3. Brain-wide error-screening system

GTree is equipped with an error-screening system for high-precision reconstruction. The system includes three components: 1) human-supervised reconstruction from sub-blocks; 2) reconstruction navigation on a brain-wide scale; and 3) online localization of differences between reconstructions of the same neuron. Using this system, we can easily identify reconstruction errors hidden in densely packed neurites and TB-sized datasets.

#### Human-supervised reconstruction from sub-blocks

Brain-wide reconstruction requires the analysis of TB-sized datasets, which are far beyond computer storage limits. Therefore, a whole-brain dataset must be divided into sub-blocks, and reconstructions from a large number of sub-blocks constitute a reconstructed neuron on a brain-wide scale. The tree-like structure of neurons means that one reconstruction error will accumulate in subsequent reconstructions. Thus, human-supervised reconstruction from each sub-block is essential for obtaining high-precision reconstructions at a brain-wide scale. This function allows the visualization of reconstructions from sub-blocks and enables us to check reconstruction errors. If there are no errors, we click the button and import the next sub-block automatically. Otherwise, we revise reconstruction errors, such as a missing neurites or miss-assigned neurites, to avoid the accumulation of errors in the subsequent imported blocks.

#### Reconstruction navigation at a brain-wide scale

As described above, a series of human-supervised reconstructions from sub-blocks constitute the brain-wide reconstruction of a neuron. The sub-block is tens of MBs in size, and the whole-brain dataset has a size of TBs. Thus, the brain-wide reconstruction of a neuron requires the analysis of hundreds of sub-blocks at a minimum. The long reconstruction process means that errors are unavoidable, despite the human response to the reconstruction. Therefore, we designed brain-wide reconstruction navigation for easily finding reconstruction errors hidden in TB-sized dataset. The tree-like structure of neurons can be mapped to a tree-graph, in which the root node represents the soma, while the nodes in the first layer represent neurites connected with the soma, and so on. Hence, by visiting the nodes of the tree-graph from the top down, the reconstructed neurites can be browsed in an orderly manner. When browsing the reconstructed neurite along with its skeleton, the neighboring image region is extracted and visualized, and all reconstructions included in the extracted region are simultaneously visualized. The checking of the reconstruction is based on the visualized information.

#### Online localization of differences between reconstructions of the same neuron

For a more challenging dataset, the general strategy for producing trustworthy results is for two or more skilled annotators to perform reconstruction of the same neuron and take the consensus results among annotators as the output reconstruction^28^. This strategy is built on fast localization of differences between reconstructions of the same neuron. A reconstructed neuron at a brain-wide scale generally contains hundreds of thousands of skeleton points. The localization of the differences between two reconstructions can essentially be performed by quantifying the differences in position between two groups of reconstructed skeleton points. We constructed a multi-dimensional Hash container^41^ for quantifying these differences rapidly in detail (**Supplementary Fig. 2**). Briefly, we assigned two groups of skeleton points to their own Hash containers and searched the matching points in these two Hash containers. Based on the search results, the locations where differences between two reconstructed skeletons appeared were labeled. Thus, the localization of differences between reconstructions of the same neuron could be achieved.

### 4. Methods for high-level automated reconstruction

The GTree software contains algorithms for reconstructing neurons, including algorithms for locating and segmenting soma, tracing neurites, and network partitioning, which are commonly used for automated reconstruction. There are also two specific algorithms integrated into GTree for this purpose. One identifies weak neurite signals, and the other detects the optimal skeleton of neurons.

#### Identification of neurites with weak signals^30^

It is a common sense that small-radius neurites will exhibit weak signals, which challenges neurite-tracing methods. Hence, we observed the characteristics of neurites and identified rules to propose a method for the identification of weak neurite signals. We found that in neuronal images, the local background was smooth, and neurites presented a strongly anisotropic shape. These image characteristics were converted into feature vectors, from which the difference between the signal and background could be displayed. A combination of these feature vectors and the machine learning method were used to construct this identification method.

#### Detection of the optimal skeleton of neurites

The reconstructed skeleton of a neurite consists of a series of sequential points generated by tracing algorithms. Detection of the neurite’s optimal skeleton is essential for post-neuron morphological analysis, such as the assessment of bifurcation numbers and lengths. Here, we consider two premises upon which to correct the positions of skeleton points: a skeleton point should present the maximum image intensity in its neighborhood region; and local smoothness should be retained at most sites in a neurite skeleton. Considering on these two premises, we designed an Lasso-based model to detect the optimal skeleton. We further extended this Lasso-based model to locate the bifurcation points of a reconstructed neuron. A bifurcation point is one terminal point of a neurite linked with another neurite. It is difficult for tracing methods to detect the position of bifurcation points where neurites are tortuous or unevenly imaged. The extended model was built on our observation that at a very small scale, the smoothness of neurites can still be satisfied. These two detection models were applied to correct the automatically reconstructed skeleton to the centerline of the neuron.

### 5. Evaluation of reconstructions

We applied high precision and recall rates to quantify the reconstructions driven by the software tools employed in this study. To calculate these two indexes, the ground-truth reconstruction is necessary. We generated the ground-truth reconstruction in GTree because GTree includes an error-screening system and achieves high-precision reconstruction more easily than semi-automatic software (**See Figs. 3 c & d**). We briefly describe how to obtain the ground-truth reconstruction and calculate the indexes for evaluating reconstructions. The pipeline for generating the ground-truth reconstruction was general. It included the following steps: 1) reconstruction of the same neuron performed by three skilled annotators; 2) import of the reconstructions into GTree to automatically locate the differences between reconstructions; and 3) rechecking of the locations of disagreements and application of the voting method to reach an agreement among annotators regarding the reconstruction of rechecked locations. Some additional notes about the above pipeline are as follows. In step 1), when reconstructing a neuron at a brain-wide scale, reconstruction navigation was employed to reduce the number of reconstruction errors. In step 2), a Hash container was employed in GTree for rapidly locating differences in the reconstructions at a brain-wide scale. In step 3), careful checking of the reconstructions, especially in 2D view mode, can confirm whether the checked reconstruction is accepted or not (**Video 10**). By analyzing the massive datasets, we found that if attention was focused on the reconstruction, errors were actually commonly avoidable. Therefore, the statistical model with stricter criteria for ground-truth reconstructions was not employed here.

By comparing the ground truth to reconstructions driven by software tools, we calculated two reconstruction evaluation indexes ^18^: the precision and recall rates. The reconstruction consists of a series of skeleton points that are located at the center of neurites and connect to each other. Hence, for a given skeleton point in the ground-truth reconstruction, we searched the point from the reconstruction to be evaluated that was nearest to the given point. If the distance of these two matched points was less than the predetermined threshold (8 μm), both the given point and the searched point were regarded as true positive points. Thus, we could label all positive points in the ground-truth reconstruction and the reconstruction to be evaluated. The recall rate refers to the ratio of the number of true positive points to the number of all skeleton points in the ground-truth reconstruction. The precision rate refers to the ratio of the number of true positive points to the number of all skeleton points in the reconstruction to be evaluated. A predetermined parameter is required to calculate these two evaluation indexes. We set this parameter as 8 μm in our application. This setting is reasonable and is explained in detail ^18^. We also note that the Diadem score^42^, a popular index for evaluating reconstructions, was not used in our analysis because in both GTree and the semi-automatic software, human-supervised reconstruction was performed and could achieve highly accurate results, leading to very high Diadem scores.

### 6. Other methodology

#### Sample preparation

The experiments were performed in accordance with the guidelines of the Experimental Animal Ethics Committee of Huazhong University of Science and Technology. We used the C57BL/6J mouse line (adult P56 male mice) for our analysis. We adopted AAV to sparsely label neurons in the cortex.

#### Data acquisition

The whole mouse brain was imaged using fluorescence micro-optical sectioning tomography microscopy^43^. In this imaging procedure, the chemical reactivation technique was employed for high-quality images. The experimental spatial resolution of this imaging system was 0.32 μm in the lateral dimension and 1 μm in the axial dimension.

#### Quantification of the time cost of reconstruction

To accurately estimate the time cost of reconstruction using GTree and semi-automatic software, we carefully selected the annotators and trained them. Two annotators performed reconstructions using semi-automatic software. The training time for these two annotators was more than 200 hours. After this long training period, the annotators were proficient in the use of the semi-automatic software. The training time for the GTree annotators was also 200 hours.

Based on these conditions, we present the statistics regarding the time cost for reconstruction in **Fig. 2g** and **Fig. 3e**. Note that if the training time were to be further increased, the reconstruction time cost would be increased negligibly.

#### Computing platform

In quantifying the time cost of the reconstructions, the annotators worked on workstations with Windows 7 or Windows 10. The CPU of the workstations was an E5 CPU. The workstations were equipped with a dedicated video card (NVidia GTX 960) and were directly connected RAID, which included the whole-brain dataset.

#### Software availability

GTree is a freely available and can be downloaded from https://github.com/artzers/AdTree, in which some test datasets and a user guide are also included.

## Acknowledgments

We thank Drs. Yunyun Han, Pavel Osten and Arun Narasimhan for providing testing datasets. We also thank the Optical Bioimaging Core Facility of WNLO-HUST for the support in data acquisition, and the Analytical and Testing Center of HUST for spectral measurements. We thank the helps from Li Jing and Su Lei in design of the software tool. We thank Li Shoucheng, Li Yingfei, Zhao Ming, Wang Hao, Chen Cheng, Zhao Yu, Huang Lu, Kong Xinyi, Li Hanying, Zhou Wuxian, Tian Tian, He Sijie, Wang Danni and Gong Yaqiong for their efforts in testing the software and in tracing neurons. This work was supported by National Natural Science Foundation of China (Grant No. 81327802, 81771913), National Program on Key Basic Research Project of China (Grant No. 2015CB7556003), the Science Fund for Creative Research Group of China (Grant No. 61721092), Science Fund for Young and Middle-aged Creative Research Group of the Universities in Hubei Province (Grant No. T201520) and Director Fund of WNLO.

